# Decoding non-human mammalian adaptive signatures of 2.3.4.4b H5N1 to assess its human adaptive potential

**DOI:** 10.1101/2024.08.26.609722

**Authors:** Ranjana Nataraj, Avinash Karkada Ashok, Ayushi Amin Dey, Sannula Kesavardhana

## Abstract

The recent panzootic 2.3.4.4b clade H5N1 infected diverse non-human mammalian species globally, showed mammal-to-mammal transmission among them and caused sporadic human infections. However, whether 2.3.4.4b H5N1 circulating in non-human mammals can establish human infections and spread among humans is unclear. Gain-of-function research restrictions preclude assessing human adapting mutations of 2.3.4.4b H5N1. Here, we tracked the evolution of 2.3.4.4b H5N1 that infected non-human mammals and evaluated their ability to gain human adaptations. The non-human mammal 2.3.4.4b H5N1 partly acquired classical human adapting mutations, which are identical to the residues of H1N1pdm09 and seasonal human H3N2 infections while showing a few species-specific adaptations that might be potential barriers for successful human adaptations. Despite minimal changes in Hemagglutinin (HA), A160T and T199I mutations near the receptor binding site of HA in dairy cattle viruses indicate the rapid HA glycan surface evolution affecting virus entry and immune evasion. The quantitative assessment indicated that 2.3.4.4b H5N1 circulating in bears, cattle, dolphins, and foxes show higher human adaptive potential than other hosts. Also, H5N1 infections in mammals across time showed a unique set of adaptations in the 2.3.4.4b clade compared to previously circulating strains, especially the acquisition of Q591 adaptation in PB2 that enables human adaptation. Thus, 2.3.4.4b H5N1 acquires human adaptations due to natural selection pressure in non-human mammals. Overall, our study delineates human adaptation and infection risk of specific non-human mammalian circulating HPAI 2.3.4.4b H5N1 strains.

GRAPHICAL ABSTRACT

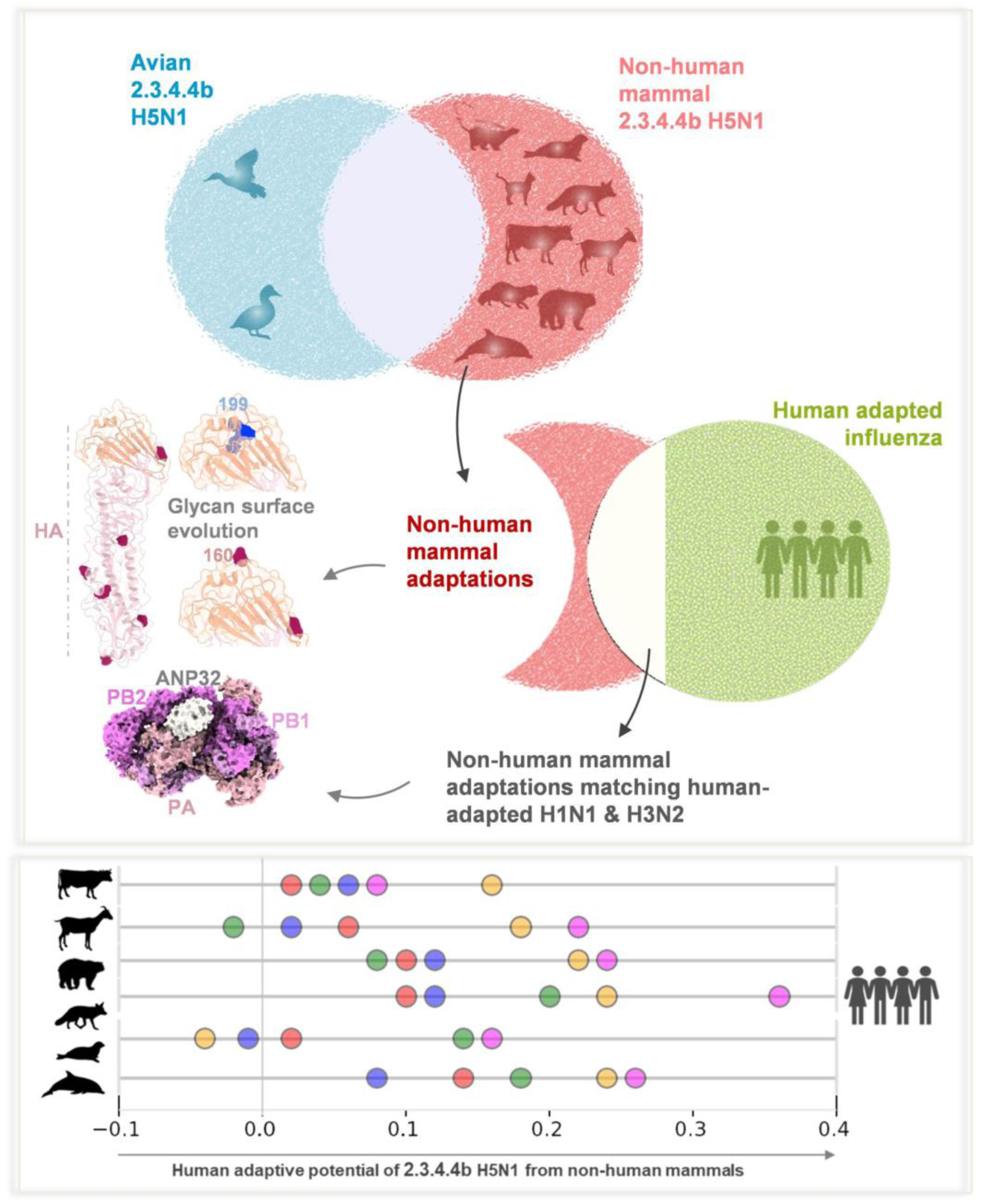

## RESULTS

Highly pathogenic avian influenza (HPAI) viruses circulate in poultry and domestic birds and often reassort with wild bird influenza subtypes. The HPAI H5N1 causes frequent outbreaks in poultry and domestic birds and is occasionally transmitted to humans. The recent 2.3.4.4b clade influenza A(H5N1) virus showed unprecedented infections and spread in diverse species^1–4^. This virus acquired the ability to spread through wild birds, which enabled transmission to various marine and terrestrial mammalian species globally^5–10^. H5N1 is adapted to replicate primarily in avian species. Its ability to replicate and spread in mammals requires specific adaptations in its surface glycoproteins, hemagglutinin (HA), and neuraminidase (NA), and polymerase complex required for virus replication and transcription. Recent studies demonstrate that 2.3.4.4b H5N1 viruses were able to spread in infected cattle^11,12^, minks^3^, sea lions^2^, and experimentally infected ferrets^11^. This may enable the emergence of novel strains suitable to replicate and propagate in mammals, including humans, thus posing a threat to public health and the economy. In this work, we monitored the evolutionary trajectories of 2.3.4.4b H5N1 in infected non-human mammals to define the mammalian-specific adaptations of this virus and its current potential in establishing human infections.

Upon transmitting to non-human mammalian hosts, the avian influenza H5N1 genome rapidly acquires mutations that enable adaptation to the new host cellular environment. To monitor the adaptation of 2.3.4.4b H5N1, avian and non-human mammal-infected virus sequences were analysed to establish the residue specificities of H5N1 proteins from avian and mammalian species. First, we studied which protein of the 2.3.4.4b H5N1 is under immediate selection pressure once introduced into non-human mammalian hosts by monitoring amino acid residue site variations in each virus protein (**Fig. 1A**). Among all the proteins of H5N1, the Non-Structural Protein 1 (NS1) was subjected to immediate selective pressure, with approximately 5% of NS1 acquiring mutations shortly after infecting a non-human mammalian host (**Fig 1A**). This is particularly intriguing given that NS1 is the smallest protein of the influenza virus and counteracts host defense mechanisms. In addition to NS1, the components of the heterotrimeric influenza polymerase complex (FluPolA), polymerase acidic (PA), polymerase basic protein 1 (PB1), and protein 2 (PB2), also showed an increased number of mutations in infected non-mammalian hosts (**Fig. 1A**). However, HA showed the fewest modifications compared to the rest of the H5N1 proteins (**Fig. 1A**). This indicates minimal HA adaptation in avian 2.3.4.4b H5N1 that established infection in non-human mammalian hosts.

**Figure 1:**
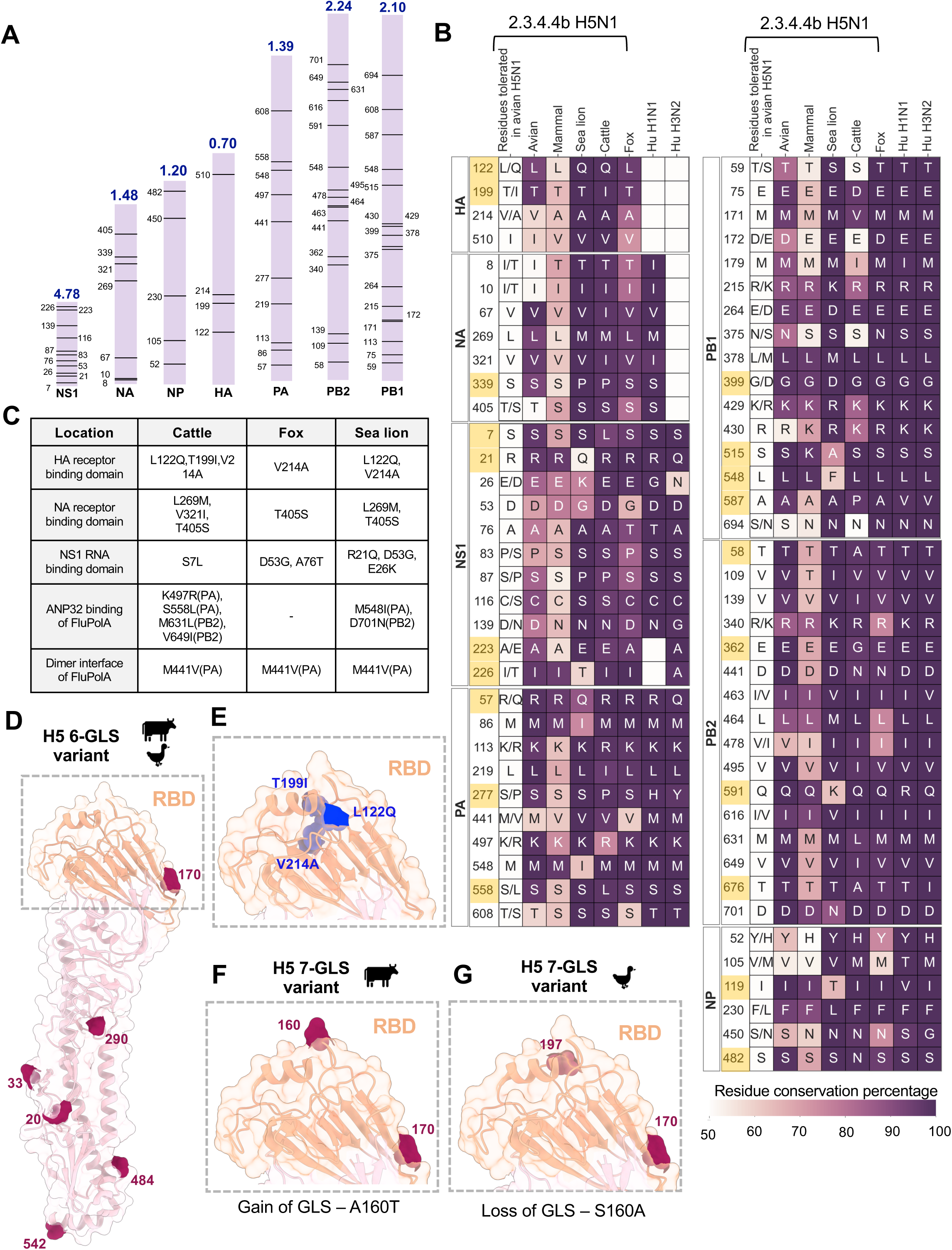
Non-human mammal-specific adaptations of 2.3.4.4b H5N1 reveal rapid evolution of glycan surface at the receptor binding site of HA. **A.** The points of variation between avian and non-human mammalian-adapted 2.3.4.4b H5N1 are shown protein-wise. The length of each bar corresponds to the size of the protein, with bars arranged in increasing order of protein size. Horizontal lines within each bar indicate amino acid positions with differing residues in avian and non-human mammalian adapted 2.3.4.4b H5N1. The total number of variable points is normalized by the size of the protein, shown above each bar. **B.** The residue identity and conservation for the positions identified in Fig 1A are depicted for 2.3.4.4b H5N1 adapted to avian, mammalian, sea lion, cattle, and fox hosts, as well as for the H1N1pdm09 and H3N2 strains adapted to humans. The first column, labeled “Residues tolerated in avian H5N1,” notes all amino acid variants at each position in the avian-adapted variant, regardless of coverage. Each cell in the heatmap is annotated with the amino acid residue circulating in at least 50% of the isolates, with the cell color representing this level of residue coverage. The residues where de novo mutations (residues not tolerated in avian H5N1) occurred are highlighted in yellow. **C.** The table summarizes the mutations from Fig 1A and B, contextualizing them within the important functional surfaces of the protein for the three host types: cattle, fox, and sea lion. **D.** The six glycosylation sites variant of HA of 2.3.4.4b H5N1 adapted to avian and dairy cattle hosts shown on a monomer of H5 from the trimer assembly (PDB:3UBE). The receptor binding domain of H5 is shown in orange while the rest of the monomer is shown in pink. Glycosylation motif positions (NxS/T) are shown as red spheres and the sites are annotated by the position of the N (Asn) residue in H3 numbering. GLS – glycosylation sites. **E**. The non-human mammalian adaptations of 2.3.4.4b H5 mapped onto the receptor binding domain of an H5 monomer (PDB:3UBE). The mutations are shown as blue spheres. The spheres are annotated to highlight the change in residue identity from avian to mammalian host. **F,G**. The glycosylation coverage variants of cattle and avian adapted 2.3.4.4b H5N1 H5 are mapped onto the RBD of an HA monomer in the H5 trimeric assembly (PDB: 3UBE). The H5 monomer RBD is represented by a light orange ribbon and surface model. Glycosylation motif positions (NxS/T) are shown as red spheres, with the position of the N residue mentioned in H3 numbering. The mutation events leading to the formation of these glycosylated variants in cattle and avian hosts are mentioned below the inset.

We further annotated the specific variations (with a minimum of 50% coverage) that appeared once the virus jumped to non-human mammals to relate whether these variations are preadaptive mutations in circulating avian H5N1 or they occur de novo (the residues different from preadaptive mutations). Due to the sequence sampling constraints, we analyzed fox, cattle, and sea lion viruses individually, and the rest of the non-human mammal viruses have been combined. These residue positions were further compared with avian 2.3.4.4b H5N1, pandemic 2009 H1N1, and seasonal H3N2 viruses. We found that the H5N1 that infected non-human mammals showed multiple de novo amino acid substitutions (**Fig. 1B**). These observations suggest the rapid adaptation of NS1 and polymerase complex proteins of 2.3.4.4b H5N1 in non-human mammals. Mappings of the mutations that the 2.3.4.4b H5N1 strain acquired upon infecting non-human mammals revealed several mutations at the functional surfaces of HA and NA (**Fig. 1C-D & Fig. S1A**), NS1 (**Fig. 1C & Fig. S1B**) and polymerase complex proteins (**Fig. 1C & Fig. S1C-D**). T199I mutation in H5 HA of the dairy cattle viruses disrupts a glycosylation motif near the receptor binding site (**Fig. 1E**). We recently reported that 2.3.4.4b H5N1 evolved to have an H5 variant with seven glycosylation sites that determine its preferential pairing with long stalk N1, a characteristic feature of 2.3.4.4b H5N1^9^. Although most of the avian 2.3.4.4b H5N1 viruses show seven glycosylation sites variant of HA, a small fraction of circulating avian 2.3.4.4b H5N1 lost a glycosylation site due to T199I mutation^13^ (glycosylation motif, NPT -> NPI) and still paired with long stalk N1^9^. Indeed, we found that one of the first avian 2.3.4.4b H5N1 isolate (EPI_ISL_19014396) transmitted to dairy cattle^14,15^ already showed T199A (instead of T1991I) mutation, suggesting the selection pressure on this residue in avian hosts and thus does not represent an adaption in dairy cattle. Intriguingly, in recent cattle infections, A160T mutation appeared in HA (17 sequences out of 932 H5 sequences of cattle), potentially restoring a glycosylation site near the sialic acid binding site in the receptor binding domain (RBD) (**Fig. 1F-G**). Loss of glycosylation at 160 residue position of avian HA in 2.3.4.4b H5N1 confers seven glycosylation site variants of H5, which is critical for pairing H5 with long stalk N1 and the functional balance required for virus fitness^9^. However, N1 sequences in cattle H5N1 viruses bearing HA with A160T mutation continued to show long stalk. Also, a recent deep mutational scanning study showed that A160T mutations in 2.3.4.4b H5 HA enable the virus to escape from neutralizing sera^16^. This suggests the evolving glycan surface of H5 HA in cattle, which potentially impacts the receptor binding affinity and immune evasion.

The FluPolA also showed significant changes in the functionally important surfaces. The replication platform of influenza consists of an asymmetric dimer of FluPolA heterotrimers, bridged by the host acidic nucleophosphoprotein of 32KDa (ANP32)^17^. Upon binding to host ANP32, the influenza FluPolA switches from transcription to replication. ANP32 appears in A and B isoforms. ANP32A is the only ANP32 family member that efficiently supports FluPolA in birds, while in humans and most other mammalian hosts, isoforms ANP32A and ANP32B support FluPolA activity to varying levels^18^. A major barrier to avian influenza virus replication in mammalian cells is the incompatibility of the FluPolA with host-specific ANP32 proteins^19^. Structure-based mapping of the non-human mammalian 2.3.4.4b H5N1 mutations of FluPolA^20^ showed that two of the residues of the PB2 subunit involved in ANP32 binding were mutated (**Fig. S1C**). These mutations (M631L in cattle and D701N in sea lions) have been previously identified to confer human adaptation of avian viruses. M631L and D701N mutations in the ANP32 binding site allow the FluPolA to utilize both ANP32A and 32B isoforms^18,21,22^. These mutations are unlike the well-characterized human adaptation E627K, which biases the FluPolA to use ANP32B proteins instead of isoform A^18^. Cell line-based studies also showed that D701N mutation facilitates increased adaptation of the avian influenza virus in human cells^18,23^. In addition, a few residues, M548, S558 and K497, in PA and V649 in PB2 of FluPolA, which are in the ANP32 binding pocket (<3.5Å distance from ANP32 binding surface), were mutated in cattle (S558L and K497R), foxes (V649I) and sea lions (M548I) (**Fig. S1C**). Analysing H1N1pdm09 and seasonal human H3N2 FluPolA for these residue positions showed that they tolerated the same amino acids as avian 2.3.4.4b H5N1 (**Fig. 1B**). Perhaps these PA residues were mutated in cattle viruses for species-specific adaptation in cattle, which might be a barrier to human adaptation.

The innate immune system restricts influenza infection at acute stages. The NS1 protein of influenza is a crucial host restriction factor to mitigate interferon and other intracellular viral restricting mechanisms^24^. NS1 of non-human mammal infected 2.3.4.4b H5N1 acquired notable variations in the RNA-binding domain of NS1. These four mutations (S7L, R21Q, E26K, D53G) in the RNA-binding domain of the NS1 are de novo and host-specific (**Fig. S1B & Fig. 1B**). This suggests the possible mammalian-specific adaptation of NS1 to facilitate vRNA replication and restricting IFN production. However, these non-human mammalian H5N1 NS1 variations were distinct from human viruses (**Fig. 1B**) and might be a potential barrier for the dairy cattle viruses to adapt to humans.

2.3.4.4b H5N1 has transmitted to more diverse mammalian species than previous isolates and showed successful spread among mammals. We analyzed all the H5N1 sequences that infected non-human mammals across time deposited in GISAID. We found that a majority of the variations were similar between 2.3.4.4b and previous H5N1 clades (**Fig. 2**). However, H5N1 acquired a few unique variations in non-human mammals with the onset of emergence of 2.3.4.4b clade H5N1 (**Fig. 2**). In particular, Q591K in PB2 appeared only in 2.3.4.4b H5N1 that was not seen in previous non-human mammal infected H5N1 viruses (**Fig. 2**). The Q591K mutation is similar to the human adapting mutation in PB2 (Q591R)^18^.

**Figure 2:**
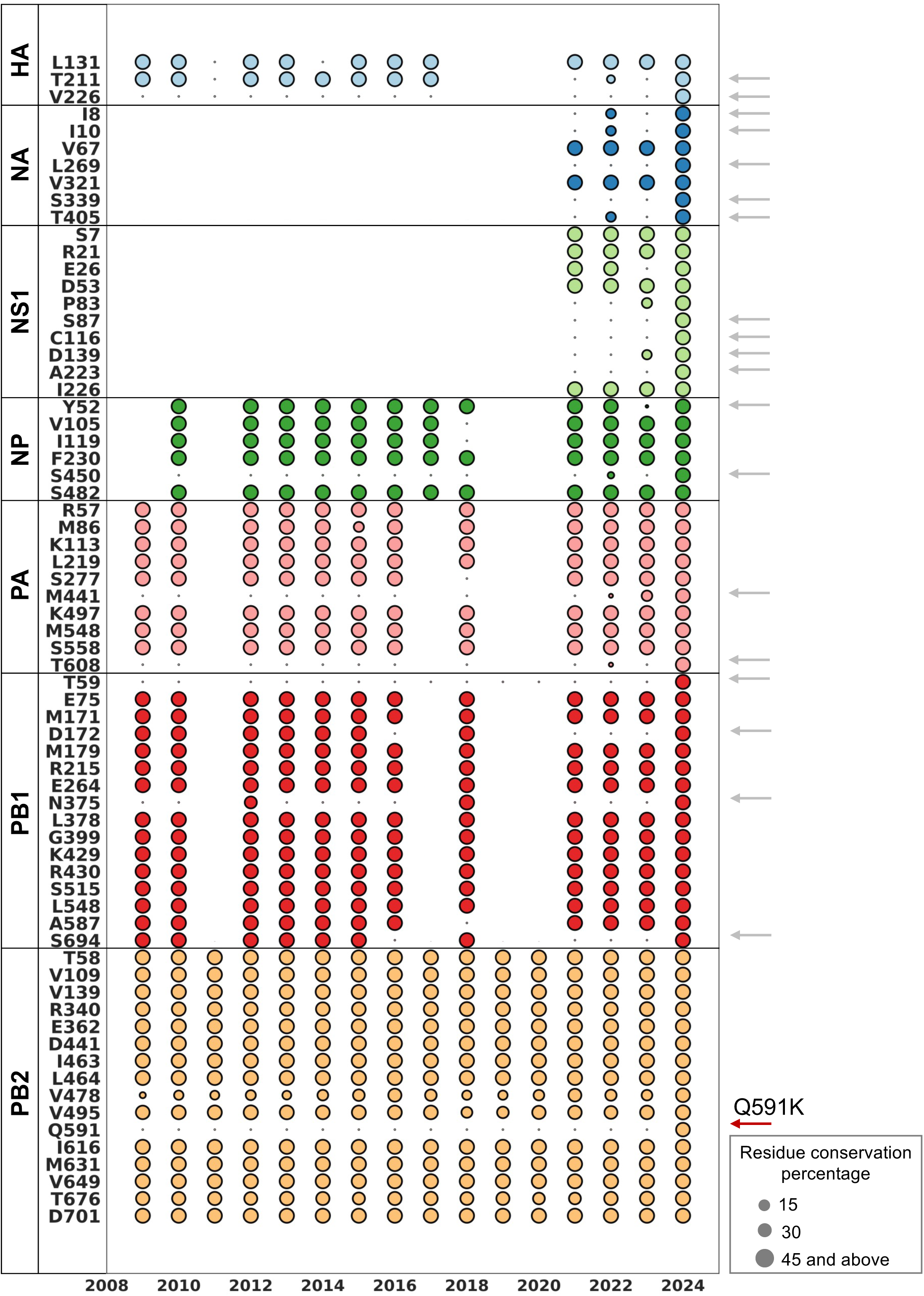
Evolutionary timeline of non-human mammal adaptations in 2.3.4.4b H5N1. The bubble plot displays the variation in the prevalence of non-human mammalian adaptations across different proteins identified in Figure 1, in mammals infected with previous clades of H5N1. The x-axis represents the years sampled, with each bubble corresponding to the number of sequences isolated from mammals infected with H5N1 during that particular year that exhibited the specific adaptation found in mammals infected with the 2.3.4.4b H5N1 clade. Bubbles are color-coded by protein type, and their size indicates the percentage coverage of the mutation. Empty spaces (without bubble representation) represent a lack of high-quality sequence information in that particular year. Arrows on the right side of the plot indicate unique adaptations acquired by the 2.3.4.4b H5N1 clade in non-human mammals. The Q591K in PB2 is a unique adaptation in 2.3.4.4b H5N1, similar to the Q591R human adaptation signature.

We then asked whether 2.3.4.4b H5N1 in non-human mammals show human permissive mutations similar to pandemic 2009 H1N1 and seasonal H3N2 viruses. To understand this, we compared the non-human mammal adapting mutations of 2.3.4.4b H5N1 (from Fig.1B) with the H1N1pdm09 and seasonal human H3N2. This analysis revealed that non-human mammal infected 2.3.4.4b H5N1 signatures (identified in Fig. 1B) converge with human-adapted residue positions of influenza in multiple viral proteins. Several seminal studies, including previous gain-of-function experiments, identified mutations that promote human adaptation of avian HPAI H5N1 (**Fig. 3A**)^18,25–46^. We further compared whether non-human mammals infected with 2.3.4.4b H5N1 viruses acquired classical human adapting mutations. The residue conservation map showed that several classical human adapting mutations have not yet appeared in 2.3.4.4b H5N1 circulating in non-human mammals (**Fig. 3B**). Intriguingly, N409S in PA, which confers human adaptation by modifying the FluPolA-ANP32 binding site^20,42^ appeared in all circulating viruses in dairy cattle, foxes, and sea lions (**Fig. 3B & Fig. 3C**). Similarly, highly conserved N383D (cattle, fox, and sea lions) and K615N (cattle) human adapting mutations were observed in PA (**Fig. 3B**), which are positioned at the FluPolA dimer interface (**Fig. 3D**). Also, a few PB2 residues crucial for ANP32 binding showed variations that match human adapting mutations. The two adapting mutations of PB2 in cattle and sea lions, M631L and D701N (**Fig. 3B**), enable human adaptation by modifying the ANP32 binding pocket. The sea lion viruses showed acquisition of Q591K (>50% coverage) in PB2 like human adapting Q591R mutation (**Fig. 3B**). These adaptations suggest the possible emergence of classical human adapting mutations in 2.3.4.4b H5N1 of non-human mammals.

**Figure 3:**
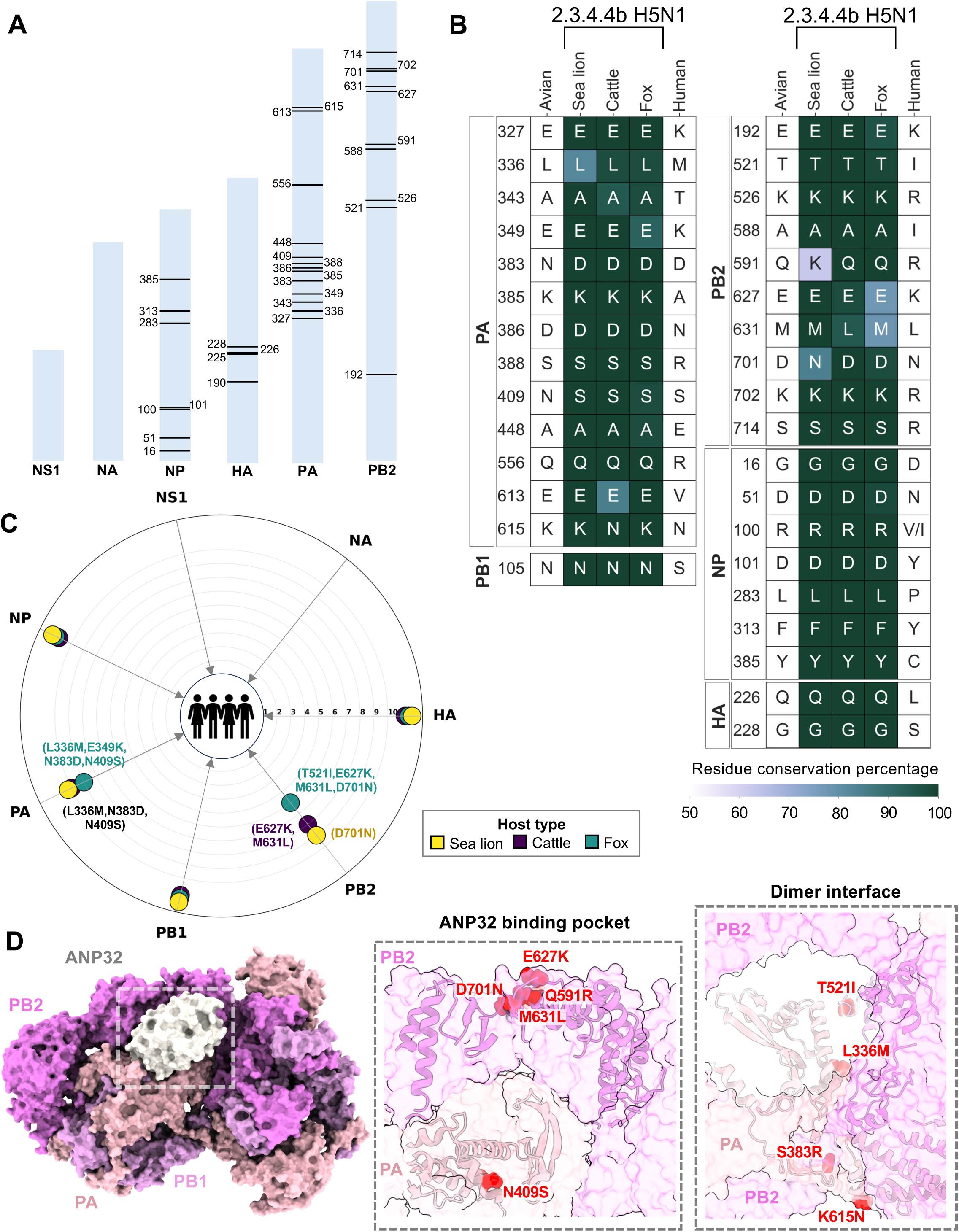
Partial gain of classical human adaptive signatures in PA and PB2 of non-human mammal-adapted 2.3.4.4b H5N1. **A.** Amino acid positions crucial for human adaptation are mapped onto the length of each protein. The bar lengths reflect the relative lengths of the proteins. For NS1 and NA, specific residues implicated in human adaptation studies were not found and are thus not annotated. **B.** Residue conservation maps show the identity and prevalence of residues at positions identified in Panel A in sea lions, cattle, and foxes infected with 2.3.4.4b H5N1. The “Human” column indicates the substitutions required at these positions to achieve human adaptation. Each cell is annotated with the residue circulating in at least 50% of the isolates, and cells are color-coded based on the exact coverage value. **C.** The radar plot illustrates all the classical human adaptive signatures that have entered circulation, regardless of the coverage of these mutations. The angular axis represents the seven H5N1 proteins investigated in this study, while the radial axis denotes the number of classical human signatures yet to be acquired in each IAV protein. The bubbles are color-coded by host type, and each bubble is annotated with the classical human signatures that have already been acquired by the virus in the corresponding host. **D.** Structural mapping of the classical human adaptive signatures already acquired by 2.3.4.4b H5N1 circulating in non-human mammals. The classical human mutations in the PA and PB2 subunits of the influenza A polymerase (FluPolA), identified in Fig 3C, are mapped onto the ANP32 binding pocket and dimeric interface of PDB:8R1J.

Intriguingly, a few specific positions of PA and PB2 proteins of H5N1 in cattle, foxes, and sea lions corresponding to human adapting positions showed reduced coverage (**Fig. 3B**), suggesting the existence of additional adaptions that have not yet dominated in circulating viruses. This prompted us to examine less frequent variations in 2.3.4.4b H5N1 of non-human mammals to assess their ability to emerge into novel variants that resemble human-adapted influenza strains. This analysis showed the appearance of additional PA (L336M and E349K in foxes) and PB2 (E627K in foxes and cattle; T521I in foxes) human adapting variations in circulating non-human mammal viruses (**Fig. 3C-D**). Considering all these human adapting mutations in non-human mammal viruses, we found that fox PA and PB2 acquired more human adapting mutations than cattle and sea lions (**Fig. 3C**). Cattle viruses tend to show better human adaptation features in PA than PB2 (**Fig 3C**). Despite relaxing the residue coverage limit to consider less frequent variations, HA and NP did not possess any adaptations suitable to humans (**Fig. 3C**).

Furthermore, we evaluated these similarities of non-human mammal 2.3.4.4b H5N1 with human viruses to monitor the human adaptation potential of these viruses. It is reasonable to justify that the greater the number of amino acid similarities of non-human mammal H5N1 viruses with human-adapted viruses, the higher the chance that the virus establishes successful human adaptation. This is based on the simple assumption that each adaptation that occurred in 2.3.4.4b H5N1 of non-human mammals, which matches human viruses, would have an equal contribution to human adaptation. However, host-specific adaptations of 2.3.4.4b H5N1 in non-human mammals that differ from human adaptations might pose a potential barrier to further human adaptation. We found that some of the residue positions that are identical in avian 2.3.4.4b H5N1 and human H1N1/H3N2 viruses were mutated to adapt for specific non-human mammalian hosts, especially for dairy cattle (such as L219I in PA of dairy cattle, Fig. 1B). We considered these constraints for quantitatively assessing the potential of a non-human mammal H5N1 for gaining human adaptation that likely poses infection risk and spread (**Fig. 4A**). We considered 2.3.4.4b H5N1 virus sequences from diverse non-human mammals for assessing human adaptive potential. Intriguingly, this analysis showed that fox 2.3.4.4b H5N1 viruses appeared to show greater human adaptive potential in most virus proteins, followed by dolphins, bears, and cattle viruses, than all the non-human mammal viruses analyzed (**Fig. 4B-C**). Also, FluPolA components, PB1, PB2, and PA, of bear, dairy cattle, dolphin, and fox sequences appeared to be acquiring human adaptive potential (**Fig. 4B-C**). Due to the widespread dairy cattle outbreaks, the circulating 2.3.4.4b H5N1 in cattle might pose a potential human infection risk if these outbreaks continue.

**Figure 4:**
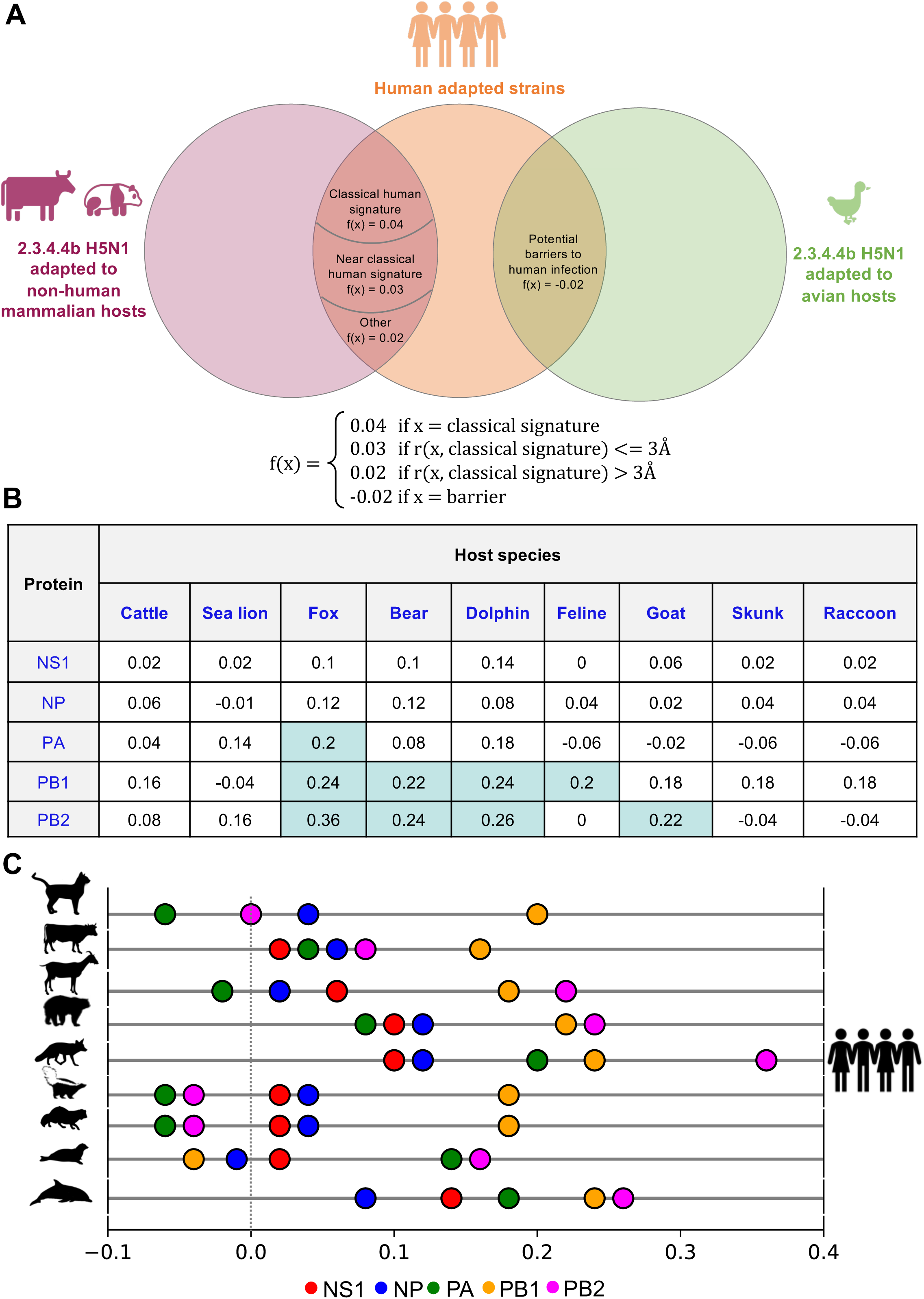
Molecular assessment of human adaptive potential in 2.3.4.4b H5N1 adapted to various non-human mammalian hosts. **A**. The Venn diagram shows the classification of adaptations in non-human mammals that mimic human influenza variants, based on their structural similarity to classical human adaptations. The diagram includes weights for each classification, assigned to each mutation x according to the function f(x). Barrier mutations, which are the mutations that are identical in both human and avian strains but have changed in non-human mammals, are also highlighted in the diagram. **B.** The human adaptive potential is calculated using the weights described in Panel A for each adaptation in influenza proteins, with the results displayed across various host types. Cells with a human adaptive potential of 0.2 or higher are highlighted in cyan. **C.** The values in Panel B for each protein are plotted on a one-dimensional graph for all host types. Each bubble is color-coded by protein type, with the x-axis representing the human adaptive potential. Negative values indicate that the barriers outweigh potential human adaptations, making the variant less potent to adapt to humans compared to the avian variant. A value of zero suggests a balance between barriers and adaptations. A positive value indicates a favorable human adaptive potential, suggesting a higher likelihood of establishing human infection compared to the avian variant.

## DISCUSSION

The recent 2.3.4.4b clade H5N1 shows unusual spread in diverse terrestrial and marine mammals and acquired mammal-to-mammal transmission ability. The sporadic human infections of circulating viruses in dairy cattle suggest the frequent spillover of this virus to humans. The increasing frequency of these infections raises concerns about potential reassortment events among influenza virus genes, which could lead to the emergence of the next human pandemic influenza A virus. In this study, we monitored 2.3.4.4b H5N1 adaptive mutations in non-human mammals to define their ability to acquire human tropism over time. We undertook comprehensive comparisons among the major encoded protein sequences of influenza and hypothesized that host-specific adaptations of 2.3.4.4b H5N1 in non-human mammals might delay successful human infections once they are transmitted. Our observations show the unique evolutionary trajectory of 2.3.4.4b H5N1 in non-human mammals compared to previous viruses. Our study provides key insights into the following critical questions.

Do non-human mammals enable 2.3.4.4b H5N1 to gain human adaptations? Several non-human mammal species acquired classical human adapting mutations, similar to the 2009 human pandemic H1N1 and H3N2. In particular, PA and PB2 of non-human mammal viruses acquired several classical human adapting mutations. These mutations in PA and PB2 expand their ability to use both isoforms of ANP32, and the 2.3.4.4b H5N1 is no longer restricted to host types with dominant ANP32B expression. This could explain why we see atypical hosts for 2.3.4.4b H5N1. This is a major concern as the range of non-human mammals that could serve as intermediate hosts for influenza viruses is expanding. We anticipate that the diverse non-human mammal host range and the ability of mammal-to-mammal transmission of 2.3.4.4b H5N1 could enable the evolution of more human adaptive mutations.

What is the selection pressure on specific proteins of 2.3.4.4b H5N1 in non-human mammals? Surprisingly, NS1 was under a stronger selection pressure than other proteins of 2.3.4.4b H5N1 viruses once they jumped from birds to non-human mammals. We found several residue positions in PA and NS1, which acquired species-specific adaptations that do not match human adaptations. These residues are identical in avian and human viruses but differ in non-human mammals. Perhaps PA and NS1 species-specific adaptations could be one of the factors contributing to the mild symptoms in humans who were infected with viruses circulating in dairy cattle. Interestingly, H5 HA shows the least adaptations in non-human mammals but rapidly changes the glycan surface near the receptor binding site. The HA glycosylation pattern and the NA stalk length likely drove the evolution of 2.3.4.4b clade H5N1, which differs from previous H5N1 isolates. The glycosylation sites involving 158 (158-160) and 197 (197-199) residues of HA have been evolving in gs/GD lineage H5 viruses and determine the major H5 variant types circulating at a given time. The dynamics of HA surface glycan patterns need to be monitored and examined if these changes bring a sudden shift in HA’s receptor binding specificity or ability to bind diverse receptors on human cells.

Which non-human mammal H5N1 is closely related to human influenza viruses and likely shows greater infection risk? This study assessed the human adaptive potential of several non-human mammals infected with 2.3.4.4b H4N1 by considering the variations in most genome segments. Interestingly, 2.3.4.4b H5N1 in bears, dolphins, and foxes show better human adaptive potential than other non-human mammals. PA, PB1, and PB2 rapidly gain human adaptations in H5N1 of non-human mammals. In particular, the adaptations of PB2 of H5N1 from foxes, dolphins, and bears show greater human adaptive potential. Dairy cattle viruses are also evolving human adaptations in polymerase complex proteins.

### Study limitations

In this study, we focused exclusively on the molecular aspects of human adaptive potential. Assuming that the likelihood of exposure to non-human mammals is constant, we evaluated the potential of 2.3.4.4b H5N1 that adapted to non-human mammals to establish infection in humans. Our findings suggest that viruses adapted to foxes and dolphins exhibit significantly greater potential for human adaptation compared to those adapted to cattle, goats, and raccoons. However, it is crucial to consider that the probability of human exposure to these wild animals is much lower than that of domesticated animals like dairy cattle and goats. While assessing adaptive potential and barriers, we recognize that mutations categorized as “barriers” may not necessarily prevent human infection, as they could rapidly mutate into human-permissive variants post-infection. The ability to fully understand these mutations is limited by restrictions on gain-of-function experiments. Also, our observations of rapid glycan surface evolution of 2.3.4.4b H5 HA need functional validation to establish the significance of these changes on receptor specificities. Nonetheless, given the evolving host range of the 2.3.4.4b H5N1 strain, monitoring even distant possibilities of human exposure remains critical.

## METHODS

### 2.3.4.4b H5N1 and human influenza datasets

A reproducible pipeline was developed to streamline the data analysis and is accessible via <u>GitHub</u>. Protein sequence datasets for HA, NA, NS1, NP, PA, PB1, and PB2 from 2.3.4.4b H5N1, sampled up until August 1, 2024, were obtained from the EpiFlu database on GISAID.org for both avian and non-human mammalian hosts (sea lions, dairy cattle, and foxes). These three host species were selected due to the constraints on sequence availability and due to their unusual roles as hosts for influenza A viruses (IAV), their diverse ecological niches, and the sustained transmission observed among these mammals. Notably, since April 2024, the Centers for Disease Control and Prevention (CDC) and the United States Department of Agriculture (USDA) have reported instances of clade 2.3.4.4b H5N1 infection in dairy cattle, and these newly identified sequences were included in our analysis. To assess the human adaptive potential of the 2.3.4.4b H5N1 variants circulating in these non-human mammalian hosts, we performed a comparative analysis using sequences of H1N1 (pdm09H1N1) and seasonal H3N2, the predominant human influenza strains known for sustained human-to-human transmission. This comparative dataset was also sourced from the EpiFlu database on GISAID.org. To ensure a comprehensive global analysis, no geographical restrictions were applied during dataset acquisition. Focusing on residue-level variability in influenza virus proteins across different host types, we retained and analyzed only full-length, high-quality sequence data. In total, 7000 sequences of 2.3.4.4b H5N1 from avian species, 820 sequences from non-human mammals (distributed among sea lions, cattle, and foxes), and 35000 sequences of human H1N1 and H3N2 were analyzed in this study.

### Protein residue conservation maps of H5N1 and human influenza viruses

To generate the residue conservation maps in Figure 1B, amino acid variations between avian and mammalian-adapted 2.3.4.4b H5N1 proteins were identified by aligning the consensus sequences of each host type. Multiple sequence alignment was performed using Clustal Omega, and consensus sequences were generated using the EMBOSS Cons package. After identifying positions under selection pressure in non-human mammalian hosts, we assessed the extent of adaptation by examining each sequence in the dataset. If a particular residue was present in at least 50% of sequences at a given position, it was used to annotate the corresponding cells in the heatmap. To normalize differences in dataset size across host types and proteins, coverage was expressed as a percentage. The intensity of each heatmap cell reflected the exact percentage of residue coverage. This analysis included all avian, non-human mammalian, and human ((pdm09H1N1 and H3N2) sequences that met quality criteria. The residue conservation maps were visualized as heatmaps using the Seaborn module in Python.

For the residue conservation map in Figure 3B, residue positions crucial for human adaptation were identified through a literature review. These positions were then analyzed for each host type, with coverage determined using the same methodology as in Figure 1B. The structures were analyzed using ChimeraX. The HA residue numbering was adapted from the Dadonaite et al. study^16^.

### Identifying non-human mammalian adaptations unique to 2.3.4.4b H5N1

The evolutionary timeline for the adaptations depicted in Figure 2 was constructed to explore whether the adaptations observed in non-human mammals infected with the 2.3.4.4b H5N1 strain are unique to this clade or represent common attributes of H5N1 infections across different clades. For this analysis, we downloaded H5N1 genomes from non-human mammalian hosts without imposing any clade restrictions. After applying quality control measures to the dataset, we obtained sequence coverage starting from 2009. For each protein, we assessed the percentage of sequences in the dataset for a given year that exhibited the adaptations uncovered in Figure 1. The coverage of these mutations was plotted year-wise as a bubble plot, with the bubble radii representing the percentage coverage for the seven proteins investigated in this study from 2009 to 2024. The bubble plot was created using the Seaborn module in Python.

### Assessment of human adaptive potential of 2.3.4.4b H5N1

A simple approach to assess human adaptive potential might involve subtracting the number of barriers from the total number of adaptations, assuming that each adaptation contributes equally to human adaptation. However, numerous studies highlight that acquiring a single classical signature, such as the E627K mutation in PB2, can significantly enhance IAV fitness in human cells. Therefore, we aimed to establish weights for each type of adaptation and barrier. To compare the molecular adaptive potential of 2.3.4.4b H5N1 strains circulating in non-human mammals with that in humans, we weighted the adaptations based on their classification as classical signatures, their proximity to these signatures, and their potential to act as barriers.

Categorizing the adaptations as:

Number of classical signatures = x

Number of adaptations within 3Å of classical signatures = y

Number of adaptations beyond 3Å of classical signatures = z

Number of potential barriers = l

The mathematical expression ax+by+cz−dl where a,b,c,d belong to the interval (0,1], was evaluated across various scenarios, including cases where barriers outnumbered adaptations (resulting in ax+by+cz−dl<0) and instances where adaptations outweighed barriers (resulting in ax+by+cz−dl>0). The weights for these mutations were empirically determined to be: a=0.04, b=0.03, c=0.02, and d=0.02. Custom scripts were developed in Python to automate the process of assigning weights to each mutation within each protein across different host types.

## ACKNOWLEDGEMENTS

We thank S.K. lab members for providing critical comments. This work was supported by funding from the Department of Biotechnology (DBT) (BT/PR45145/COT/142/24/2022), Science and Engineering Research Board (SERB-DST) (EEQ/2021/000274), Indian Council of Medical Research (ICMR) (EMDR/SG/9/2023-2019), Scheme for Transformational and Advanced Research in Sciences (STARS)-MoE grant (MoE-STARS/STARS-2/2023-0464) and the research support from thee Indian Institute of Science (IISc).

## AUTHOR CONTRIBUTIONS

S.K. conceptualized the study; R.N. performed the large sequence data sets analysis, residue conservation, protein structure analysis and adaptive variations annotations. R.N. and A.K.A. performed non-human mammal sequence analysis; R.N. and A.A.D were involved in quantitatively assessing adaptive variations. S.K. and R.N. analysed and investigated the study; S.K., and R.N., wrote the manuscript and S.K., R.N. and A.A.D. edited the manuscript; S.K. supervised the study; S.K. provided the guidance and brought the funding.

---

**Figure S1:**
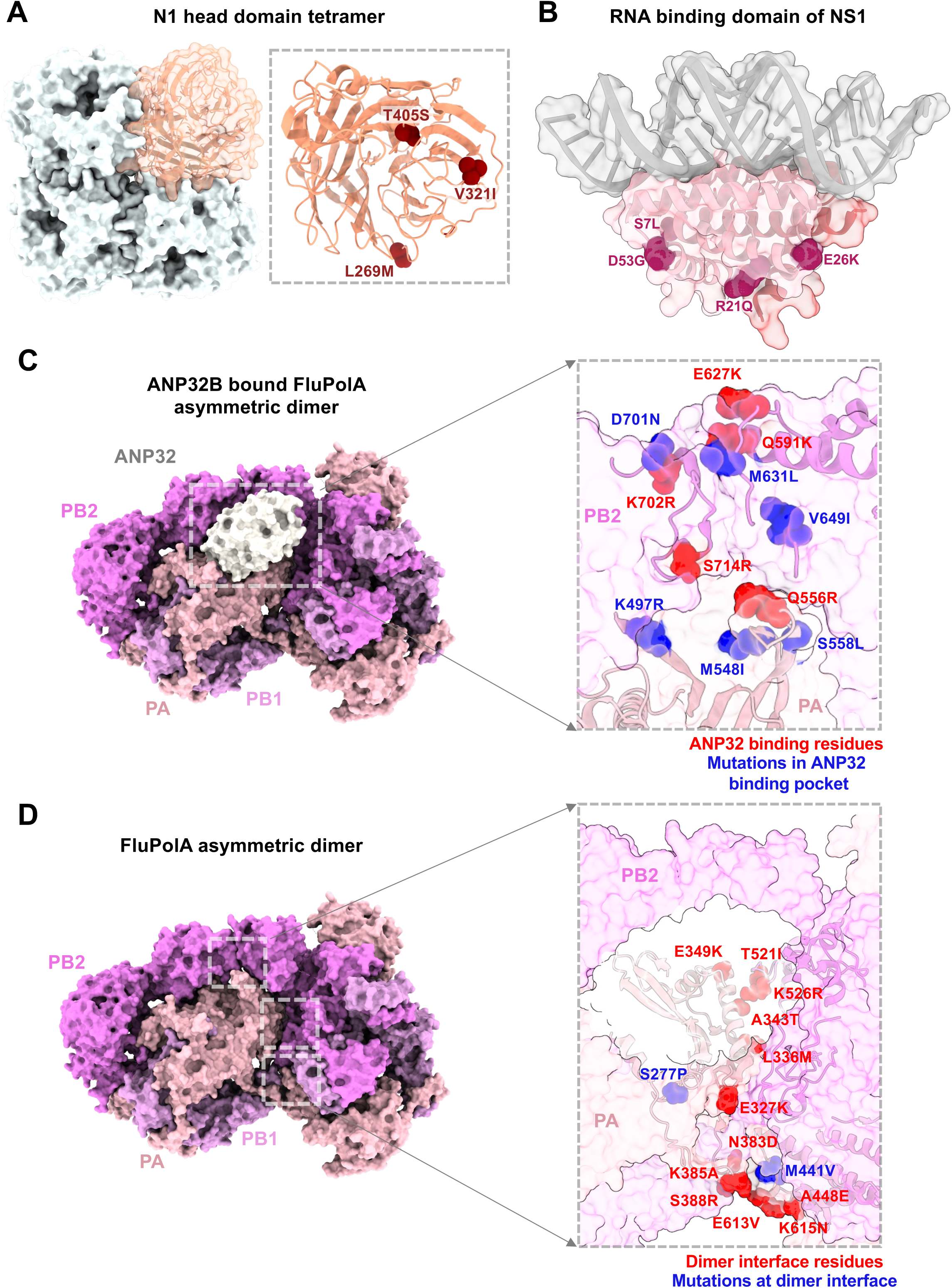
Structural mapping of non-human mammal adaptations of 2.3.4.4b H5N1. **A.** The surface representation of the N1 tetrameric head is shown (PDB:2HTY), with one monomer highlighted in orange. Zoomed-in insets show the receptor-binding domain of the N1 monomer, along with the mutations as red spheres in this domain. The positions are annotated to highlight the mutation in non-human mammals from the avian variant. **B.** The surface representation of NS1 dimer bound to dsRNA (in grey) (PDB: 2ZKO) is shown. The non-human mammalian adaptations in NS1, identified in Figure 1, are marked as magenta spheres on one of the monomers. **C.** The surface representation of the ANP32B-bound Influenza A polymerase (FluPolA) asymmetric dimer is shown, with ANP32B highlighted in white and the remaining subunits annotated (PDB: 8R1J). The zoomed-in inset reveals mutations identified in Figure-1B located within the ANP32 binding pocket of FluPolA, depicted as blue spheres. Residues directly implicated in ANP32 binding are shown as red spheres. **D.** The surface representation of the FluPolA asymmetric dimer is shown, with the subunits—PA, PB1, and PB2—annotated (PDB: 8R1J, chain G of ANP32B hidden). The zoomed-in inset highlights the mutations from Fig 1B that occur at the dimeric interface (blue), and the residues required for the interface contacts (red).

